# A novel method for tracking nitrogen kinetics *in vivo* and *ex vivo* using radioactive nitrogen-13 gas and Positron Emission Tomography

**DOI:** 10.1101/2023.06.01.543280

**Authors:** Edward T Ashworth, Ryotaro Ogawa, David Vera, Peter Lindholm

**Affiliations:** Department of Emergency Medicine, University of California San Diego, LaJolla, CA; Department of Radiology, University of California San Diego, LaJolla, CA

**Keywords:** Hyperbaric Medicine, Radiolabeling, Diving, Decompression sickness

## Abstract

**Rationale:** Decompression sickness (DCS) is caused by gaseous nitrogen dissolved in tissues forming bubbles during decompression. To date no method exists to identify nitrogen within tissues, but with advances in PET technology it may be possible to track gaseous radionuclides into tissues. We aimed to develop a method to track nitrogen movement *in vivo* that could then be used to further our understanding of DCS using nitrogen-13 (^13^N_2_) – a radioactive isotope of nitrogen that emits β+ radiation.

**Methods:** A single anesthetized and ventilated Sprague Dawley rat lay supine inside a PET scanner for 30 min. The rat breathed oxygen for the first 2 min, then was switched to a bag containing ^13^N_2_ gas mixed with oxygen for 20 min, then breathed oxygen alone for the final 8 min. Gas samples were drawn from the inspiratory line at 5, 15 and 25 min. The PET scanner recorded ^13^N_2_ with energy windows of 250-750 keV. Following the scan, a mixed blood sample was taken from the heart, while the brain, liver, femur and thigh muscle were removed to determine organ radioactivity using a gamma counter.

**Results:** The gas samples at 5 (5.7 kbq.ml^-1^) and 15 min (5.3 kbq.ml^-1^) showed radioactivity in the inspired gas that was absent at 25 min (0.1 kbq.ml^-1^), when the ^13^N_2_ was stopped. The signal intensity in the PET scanner increased from baseline (0.03) to 2-12 min (0.68±0.31), and 12-22 min (0.88±0.06), before reducing slightly from 22-30 min (0.61±0.04). All organs had radioactivity when measured in the gamma counter, with the highest counts in the liver (12593 counts.min^-1^.g^-1^) and the lowest in the muscle (2687 counts.min^-1^.g^-1^).

**Principal Conclusions:** This study successfully demonstrated a quantitative 3D imaging method of tracking nitrogen gas through the body both *in vivo* and *ex vivo* using PET.

## Introduction

During respiration gases are taken up by the body and distributed to various tissues. While gas exchange usually refers to oxygen and carbon dioxide the majority of respired air is nitrogen. Nitrogen gas is taken up by blood and is transported throughout the body where it dissolves into tissue. The amount of nitrogen taken up by each tissue is proportional to the partial pressure difference of nitrogen between the blood and the tissue, the tissue perfusion, and the tissue solubility. When each tissue is saturated with nitrogen there exists a homeostatic equilibrium in which no net flux of nitrogen occurs. As nitrogen gas is inert it minimally impacts physiological function. However, when the partial pressure of nitrogen increases, such as when SCUBA diving to depth, tissues can become more saturated than at sea-level (Doolette & Mitchell, 2001). As a result, when decompressing, such as ascent from a dive, rapid ascent to altitude, or during spacewalks, the tissues become supersaturated as the partial pressure of nitrogen exceeds the ambient partial pressure (Mitchell et al., 2022). When this happens bubbles can form as the gas expands, which can cause complications (Junes et al., 2022). Bubble formation leads to a range of symptoms including musculoskeletal pain, neurological impairment, and respiratory distress; collectively known as decompression sickness (DCS; (Mitchell et al., 2022)), which poses a severe occupational risk to commercial and scientific divers (Dardeau et al., 2012), as well as astronauts (Conkin et al., 2017). Despite this, knowledge of how DCS manifests is largely unknown, and is a product of retrospective and empirical analysis conducted after individuals are diagnosed with DCS. Methods to track the movement of nitrogen are scarce, but vital to understanding how DCS forms, varies between individuals, and how it can be treated.

One method that may provide a solution is the radionuclide nitrogen-13 (^13^N_2_). ^13^N_2_ is an isotope of nitrogen that emits beta-radiation with a half-life of 9.965 minutes. Beta-radiation can be tracked with positron-emission tomography (PET) to provide localized information as to where the radiation is coming from, essentially allowing the tracking of nitrogen (Berger, 2003). Furthermore, the emitted positron rapidly annihilates with an electron, leading to the production of two gamma rays (Richardson, 1938). These gamma rays can be measured using a gamma counter which measure smaller quantities of radioactivity in smaller volumes (Wilde & Ottewell, 1980). Several studies have previously looked at ^13^N_2_ in humans, primarily injected as part of a saline solution to investigate lung function (Vidal Melo et al., 2003; Winkler et al., 2022). One study looked at humans breathing ^13^N_2_ gas in normobaric conditions with a gamma detector placed by the knee to track nitrogen kinetics with a diving population in mind (Weathersby et al., 1986). Since this lone experiment technical developments have produced new imaging modalities, such as PET scanning, which can provide spatial and temporal resolution of radioactive substances within the body thereby increasing the range of possibilities for tracking ^13^N_2_ movement through the body.

However, the development of such a method can be challenging as the relatively short half-life that leaves a limited window in which to conduct experiments. Therefore, we aimed to develop an experimental model that could spatially and temporally track the movement of nitrogen through a ventilated rodent in a safe and repeatable manner.

## Methodology

The ^13^N_2_ was created off-site (PETNET, Siemens Medical Solutions, San Diego, CA) using a cyclotron in accordance with prior studies (Iwata et al., 1978). A liquid target containing aqueous NH_4_Cl solution (1.0 M, pH = 11) was irradiated with 15 MeV protons for 30 min. ^13^N_2_ was extracted from the target by a helium sweep gas, which passed through a P_2_O_5_ absorber to purify the gas of NH_3_ and water vapor. The products are released into a, placed inside a lead ingot and casing, delivered to the laboratory (transit time ~10 min).

### Analysis of ^13^N_2_

To confirm the presence of ^13^N_2_ a number of investigations were conducted both using the liquid that had ^13^N_2_ suspended within it, and the gas containing ^13^N_2_. Both the liquid ^13^N_2_ solution and a gaseous ^13^N_2_ sample were placed in a dose calibrator (CRC-15W, Capintec, NJ, USA). Recordings were made every minute for 20 min. Samples of the liquid and gaseous ^13^N_2_ were then placed in a gamma counter (2480 Wizard^2^, Perkin Elmer, MA, USA) to obtain their radioactive spectroscopy.

### *In vivo* experiment

Upon arrival at the laboratory the vial was placed into a dose calibrator and radioactivity recorded. The vial was then shaken for 10 s and bubbled (to release the ^13^N_2_ gas from solution) at 0.5 L.min^-1^ directly from the isoflurane vaporizer (VS1482, Visual Sonics, Canada) into a non-diffusing bag (Figure 1 – Gaseous ^13^N_2_ Bag; 112110, Hans Rudolph Inc, KS, USA) pre-filled with 7 L of oxygen-isoflurane mix (3% isoflurane). Bubbling continued for 1.5 min, causing flow of an additional 1.5 L of air into the bag, providing a total of ~8.5 L which was estimated to be required for the 30 min scan time.

**Figure 1.**
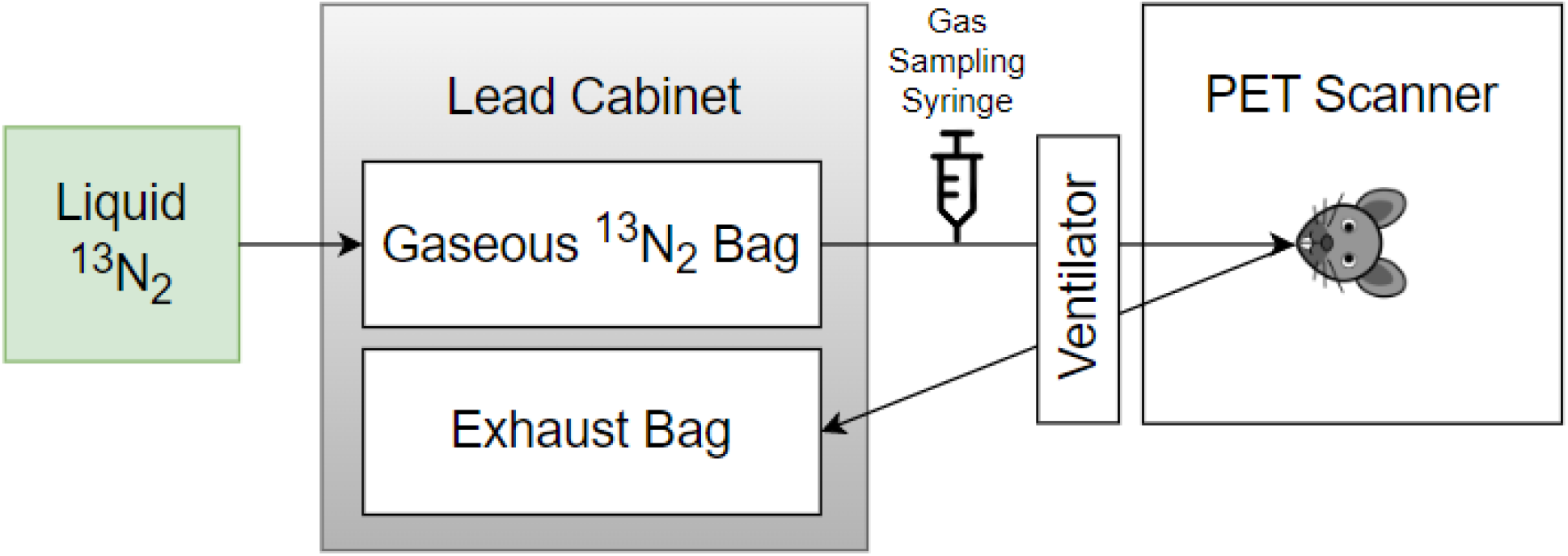
Schematic detailing how the process required to ventilate a rat with ^13^N_2_. The ^13^N_2_ arrives in liquid form and is bubbled into gaseous form which is collected in a non-diffusing bag housed within a lead cabinet. Gas is then drawn from this bag by a ventilator to a rat located inside a PET scanner, with the expired gas being collected in an exhaust bag within the lead cabinet.

A Sprague-Dawley rat (6 months, 320 g) was anaesthetized using 5% isoflurane and placed on an intubation rack (Kent Scientific, CT, USA) in accordance with institutional review board approval (IACUC, UCSD, protocol S19154). With the airway exposed an intubation tube (16G Safelet Catheter, Exel, CA, USA) with an intubation safety wedge (Kent Scientific, CT, USA) was placed down the trachea. The intubation line was then connected to a mechanical ventilator (PhysioSuite, Kent Scientific, CT, USA) set to deliver air at a rate of 90 breaths per minute, at a tidal volume of 1% body mass (i.e., 300 g = 3 ml). Isoflurane was then reduced to 3% and the rodent was set on the PET scanner (eXplore VISTA DR, GE Healthcare, IL, USA) gantry, with the center of the PET located at the level of the lungs. The PET scanner recorded ^13^N_2_ using a dynamic emission scan for 30 min with energy windows of 250-750 keV. During the first 2 min of the PET scan the rat breathed oxygen, before being switched to ^13^N_2_ at 2 min. After 22 min of scanning had elapsed the inspiration line was switched back to the oxygen.

After 5 min, and thereafter at 10 min intervals, a gas sample was obtained from the inspiration line connected to the ventilator using a 10 ml gas syringe to determine the dose being delivered to the animal (Fig 1). The syringe was immediately placed in the dose calibrator, recorded, and then flushed into an extraction vent. Immediately prior to the gas sample collection a background radiation measurement was taken to be subtracted from the recorded values. These values were then corrected for the ^13^N_2_ half-life using Eq. 1.

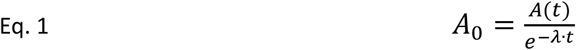

Where *A*_0_ is baseline counts per minute, *A*(*t*) is counts per minute at the time point (*t*), and *λ* is equal to 0.693/9.965 min, where 0.693 is equal to the natural logarithm of 2, and 9.965 min is the half-life of ^13^N_2_.

Upon cessation of the PET scan the rodent was surgically opened and euthanized by a mixed blood draw from the heart. Immediately following this, the liver, brain, femur and quadriceps muscle were surgically removed and, alongside the remaining rat, were placed in the dose calibrator to assess whole-rat radioactivity. The organs were then placed into a gamma counter (Gamma 8000, Beckman, IN, USA) to obtain organ-specific counts. All counts were obtained within a window of 400-600 keV. Counting continued until the uncertainty reached 2 sigma %, or 10 min had elapsed. Organ counts were corrected firstly by subtracting background radiation, and then using Eq. 1 to account for the difference in time between each sample, effectively reporting the counts when the counting process began. All organs were then weighed (CP64, Sartorius, Germany) to enable calculation of counts relative to mass.

The PET image was analyzed in Fiji image analysis software (Schindelin et al., 2012) in composite images of 2 min each. Images were converted into 3D stacks, and the entire image in each view had signal intensity recorded. Mean and standard deviation (SD) were calculated for each 2 min block. Additionally, the PET images for minutes 2-22 were amalgamated into one image to visualize the ^13^N_2_ activity in the lung.

## Results

The vial produced contained 1158.1 Mbq, which had reduced to 577.2 Mbq by the time it was measured in the laboratory. Analysis of the ^13^N_2_ liquid showed the radioactive decay was less than expected (Fig 2A), however, analysis of the ^13^N_2_ gas showed decay as expected, with a slight increase in error when measuring smaller radioactivity quantities (Fig 2B).

**Figure 2.**
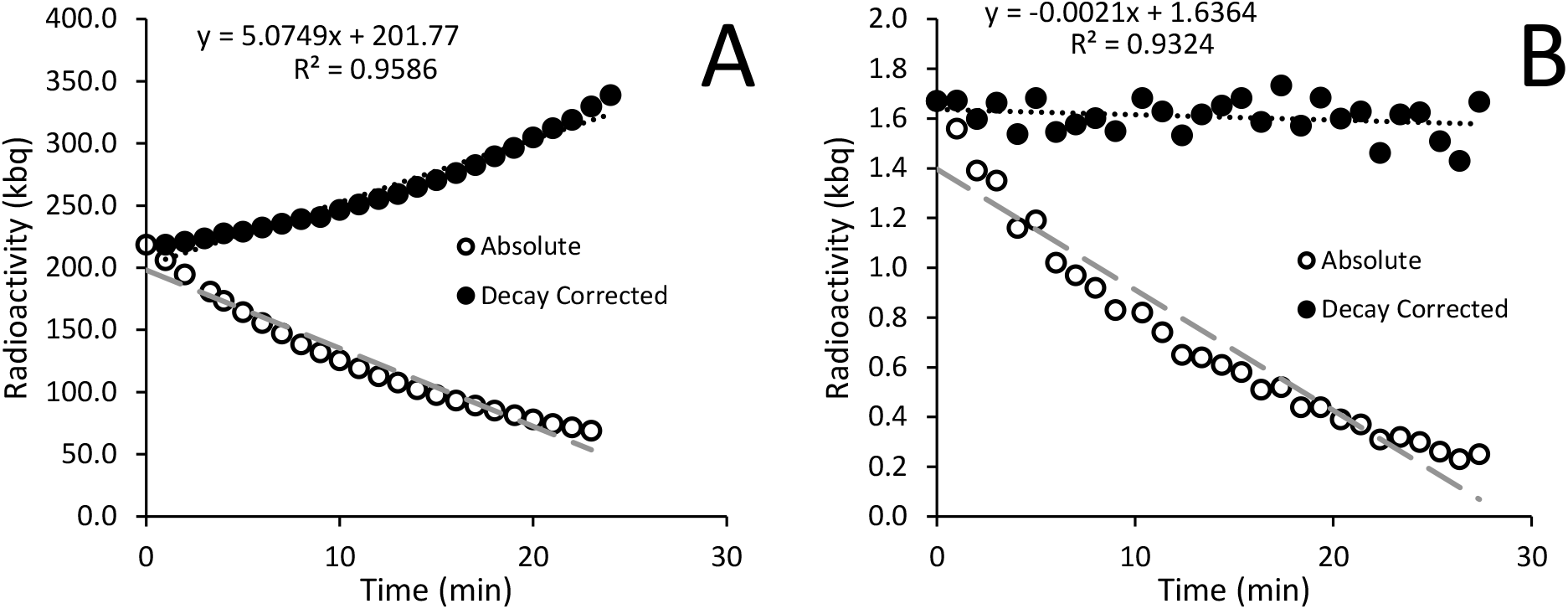
Radioactivity in the ^13^N_2_ liquid (A) and gas (B) recorded every minute. Results are displayed as both absolute values (open circles) and values adjusted for ^13^N_2_ radioactive decay (closed circles). The increase in decay corrected activity in Panel A suggests the liquid contains something other than ^13^N_2_, which was no observed in Panel B, which contains only gas.

The bubbling of the vial contents left 418.1 Mbq in the vial, indicating that ~75.1 Mbq of the original dose (when corrected) had been removed from the vial as ^13^N_2_ and entered the gaseous ^13^N_2_ bag. The in-line gas concentration was consistent between the two measurement periods, and was reduced when the ^13^N_2_ flow stopped (Table 1).

**Table 1.**
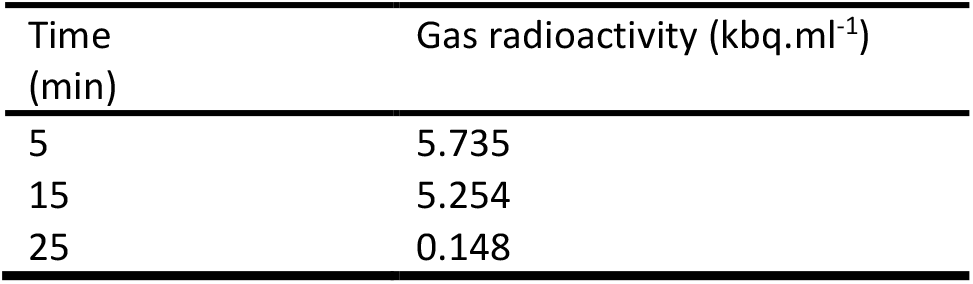
Radioactivity of inspiration line gas samples while breathing ^13^N_2_ gas. Samples are corrected for radioactive decay. ^13^N_2_ gas was turned off after 22 min.

| Time (min) | Gas radioactivity (kbq.ml<sup>-1</sup>) |
| --- | --- |
| 5 | 5.735 |
| 15 | 5.254 |
| 25 | 0.148 |

The signal intensity recorded by the PET increased once the ^13^N_2_ was connected and approached a plateau before reducing in the final minutes of the experiment (Fig 3). Visual analysis of the PET image reconstruction clearly shows the shape of the lungs being filled with ^13^N_2_, seen alongside the intubation tube and trachea which show the highest concentrations of positron emission (Fig 4).

**Figure 3.**
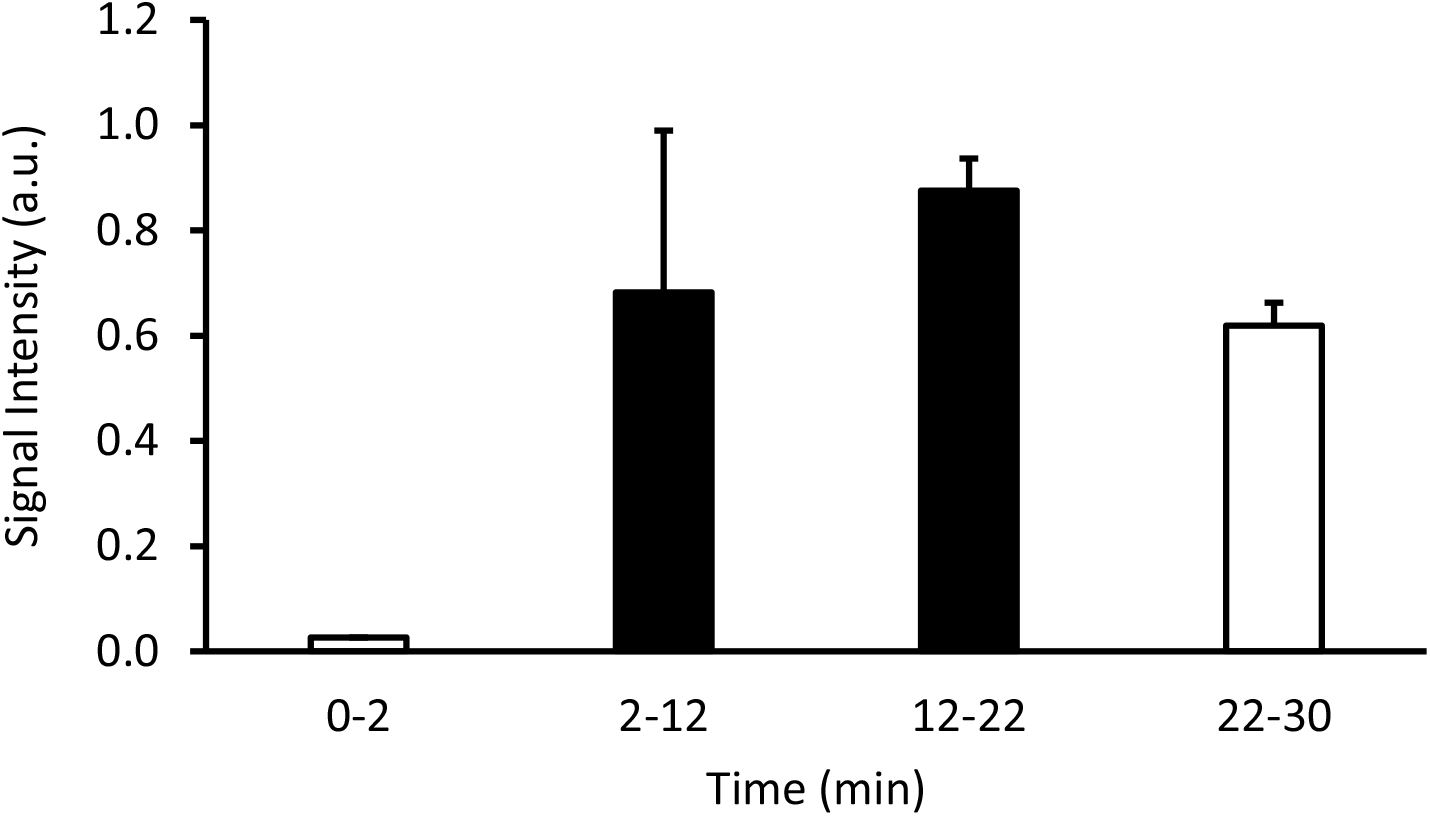
Signal intensity in arbitrary units (a.u.) over time showing the changes in radioactive positron emission from a sedated rodent breathing ^13^N_2_ gas. Values shown are an average signal intensity of the specified interval. An oxygen-isoflurane mix was breathed from 0-2 and 22-30 min (white bars), while the ^13^N_2_ gas was turned on from 2-22 min (black bars).

**Figure 4.**
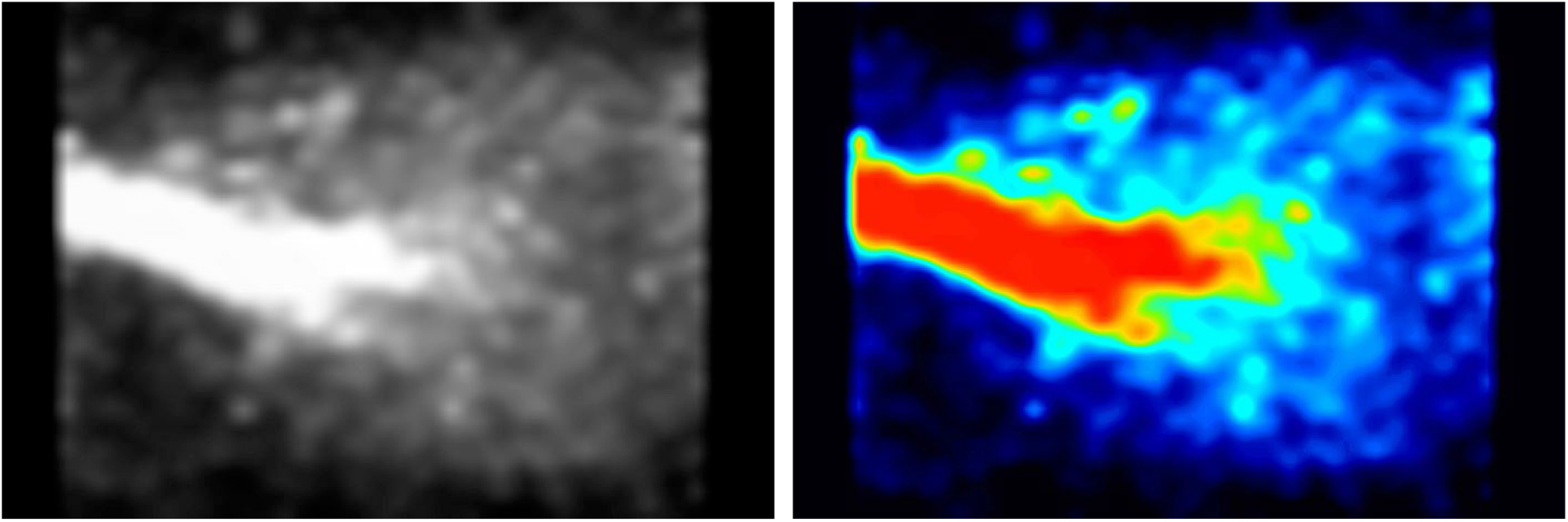
PET image of rat lung while breathing ^13^N_2_ gas. Recording window was set to 250-750 keV. The trachea conducts all of the radioactive gas, and therefore has the strongest signal (red). The radioactivity then spreads throughout the lungs, being more condensed within the middle of the lungs as the lungs are continuously moving due to respiration.

Upon conclusion of the PET scan the whole rat had 58.1 kbq of radioactivity. The liver, brain and bone had higher relative counts per minute than the blood, whereas the muscle had lower counts (Table 2).

**Table 2.** Organ parameters following breathing ^13^N_2_ for 20 min followed by 10 min of breathing oxygen. All counts were corrected for radioactive decay (Eq. 1).

| Sample | Mass (g) | Error (%) | Counts.min<sup>-1</sup>.g<sup>-1</sup> |
| --- | --- | --- | --- |
| Blood | 3.722 | 1.99 | 3390.28 |
| Liver | 8.751 | 1.91 | 12593.29 |
| Brain | 1.811 | 1.99 | 5382.77 |
| Muscle | 1.364 | 1.99 | 2686.57 |
| Bone | 0.180 | 4.53 | 3882.01 |

## Discussion

This study successfully demonstrated a method of tracking nitrogen gas through the body that could be imaged both *in vivo* using PET, where ^13^N_2_ could be tracked going in and out of the lung, and *ex vivo* using a gamma counter to assess individual organs.

The vial containing ^13^N_2_ suspended in liquid may have some radioactive contaminants (Fig 2A) but it appears that these are not bubbled out of solution and that the only radioactive component of the gas is ^13^N_2_ (Fig 2B). Furthermore, the strong linear correlation of the gaseous ^13^N_2_ decay occurred at the same rate as the half-life of ^13^N_2_ (Fig 2B), strongly suggesting that no other isotopes were present. Other isotopes that theoretically could have contaminated the liquid ^13^N_2_ include isotopes of oxygen (which have a much shorter half-life than ^13^N_2_), whereas nitrogen oxides have a much longer half-life. Therefore, it is highly likely that the only radioactive substance in the gas was ^13^N_2_.

The total radioactive volume received was only 577.2 Mbq, a safe experimental quantity, that comfortably lasted the duration of the experiment (~1 h). Future experiments could likely be of longer duration or increase the starting dose. The supply of ^13^N_2_ to the rat measured in the inspiratory line was reasonably consistent between the two measurements taken during ^13^N_2_ delivery (Table 1) suggesting good mixing of gas within the gaseous ^13^N_2_ bag. The constant supply of ^13^N_2_ increases the potential for uptake over time as only a small volume of ^13^N_2_ will be taken up with each breath, whereas if a bolus of ^13^N_2_ was delivered a large proportion could be expired.

The PET image shows ^13^N_2_ in the main areas we would expect to see it – the trachea and the lungs, with minor signal coming from the surrounding areas (Fig 4). During the first 2 min of the experiment while oxygen-isoflurane was breathed PET signal resembled background levels (Fig 3). Upon switching to ^13^N_2_ the PET signal increased steadily from 2-12 min, reaching a plateau from 12-22 min, suggesting equilibrium of pulmonary ^13^N_2_ was reached rapidly (Fig 3). Then when ^13^N_2_ delivery ceased we saw a drop in PET signal (Fig 3), effectively showing the slow washout of ^13^N_2_ from the lung and surrounding tissue. Due to the natural movement of the lungs during image acquisition it is hard to quantify the amount of ^13^N_2_ taken up in pulmonary tissue or adjacent tissue, but the subsequent organ counts showed that ^13^N_2_ uptake was widespread.

The organ counts showed higher relative ^13^N_2_ content in the liver, brain and bone than the blood (Table 2). The brain has the highest proportion of fat of any organ and is highly perfused, so might be expected to take up ^13^N_2_ more than the other organs (O’Brien & Sampson, 1965). However, these factors also mean that it will off-gas faster during the oxygen breathing period of the experiment. Indeed, neurological decompression sickness is largely reported during short dives (Lang et al., 2013; Schipke & Tetzlaff, 2016), suggestive that brain is a tissue with fast nitrogen kinetics. The liver is also highly perfused, and is the fattiest abdominal organ, which likely accounts for the large values seen here (Sijens et al., 2010). However, the liver is likely a slower tissue than the brain, as most decompression injury events involving the liver occur after more moderate dive times (Jinno et al., 2018; Siaffa et al., 2019). This means that the off-gassing of the tissue takes longer, resulting in a greater relative amount of nitrogen within the liver. Bone ^13^N_2_ content was also elevated compared to blood. Bone marrow makes up 10% of adult human total fat content which may help with nitrogen uptake. Previously Campbell and Hill (1931) assessed nitrogen composition in a variety of animals using a method utilizing Geissler’s mercury gas pump and Haldane’s apparatus. Both bone marrow and the brain were assessed, with bone marrow containing ~5 times more nitrogen than the brain. In the current study “bone” included cortical bone and marrow so marrow content alone was not evaluated, so direct comparisons cannot be drawn. A review of solubility coefficients by Weathersby and Homer (1980) showed brain and blood to have similar levels of nitrogen, with a range of approximately 50-150%. In the current study the brain has 159% more ^13^N_2_ than the blood, falling just outside of this range. Only one study used the same method to measure brain and blood nitrogen content, finding brain tissue to be 13% more soluble than blood in rabbits (Ohta et al., 1979). It is unclear whether using a perfused brain, as in the current study, would lead to similar results, and to what degree organ perfusion can be quantified.

Muscle has less fat, with human intramuscular thigh fat as low as 8% of the tissue, and therefore would be expected to store less ^13^N_2_ (Goodpaster et al., 2000). As muscle stored less ^13^N_2_ than was measured in blood (Table 2), it may have a limited capacity to take up more ^13^N_2_ as otherwise the arterial to tissue pressure gradient would continue to favor tissue uptake.

### Methodological limitations

The use of a PET scanner alone makes tracking the exact organs taking up ^13^N_2_ difficult as organ positions vary between individuals, the PET provides no anatomical landmarks, and the spatial resolution of PET is low (Vaquero & Kinahan, 2015). For the lungs which receive a large dose, their outline is easily observed due to the contrast with adjacent tissue (Fig 4), but this is not the case for other tissues. It is possible that future studies could look to use PET and computed tomography in concert to guide locating where ^13^N_2_ is, particularly in studies where survival is prioritized (Vaquero & Kinahan, 2015). In non-survival experiments, the use of the gamma counter provides a unique opportunity to gather rich data.

The 10 min half-life of ^13^N_2_ means that experiments much be short (~100 min) as otherwise the radioactive signal dissipates to that of background radiation. This severely limits the scope of experiments that can be done with this method while keeping radiation within safe limits. Decompression times during diving frequently exceed this 100 min constraint, therefore results obtained using the current method would need to be extrapolated to provide insight for these conditions. It might be possible to replenish the ^13^N_2_ supply, but this would require recalibration and multiple logistical challenges.

Usually a standardized uptake value is used to quantify PET signal (Kinahan & Fletcher, 2010). However, most studies that employ this technique use non-gaseous radionuclides that allow easy assessment of a known concentration giving a known signal. Due to the rapid decay we do not know the amount of ^13^N_2_ present in the PET and the amount of time it takes to measure takes up experimental time required for valid results to be acquired. Therefore, the method is currently limited to relative changes observed within a specific reason.

### Future applications

The current experiment was performed on a benchtop, without the compression and subsequent decompression that occurs with diving, which is where understand nitrogen kinetics would be of greater importance. It is feasible that the radioactive nitrogen could be placed inside a compression chamber prior to pressurization using the method reported here, before being compressed to the desired depth using compressed air. Alternatively, the use of a compressor could be used, but may present further logistical challenges, especially considering the short half-life. Ultimately, further development of this method will increase our understanding of DCS and how to prevent and treat it.

While this experimental paradigm has been devised for rodents and will be further developed in rodents in the future, it is hoped to be safely extended to humans in the future. Prior to this, experiments with larger animals, such as pigs will be required to ensure the radiation dose is sufficient for meaningful experiments, whilst remaining at an ethical limit.

### Disclosure

This project was supported by a grant from the US Department of Defense; ONR grant number: N00014-20-1-276

### Key points

#### Question

This study investigated a new method to determine whether nitrogen could be tracked through the body using a radioactive tracer (^13^N_2_).

#### Pertinent findings

The method deployed successfully showed ^13^N_2_ being inhaled at a level that could be detected with a PET scanner, while organs removed revealed uptake of ^13^N_2_.

#### Implications for patient care

These results lay the foundation to use this experimental design in future studies to understand how nitrogen moves through the body, especially during compression to help treat and prevent decompression sickness.

